# Mutated SF3B1 is associated with transcript isoform changes of the genes UQCC and RPL31 both in CLLs and uveal melanomas

**DOI:** 10.1101/000992

**Authors:** Alejandro Reyes, Carolin Blume, Vincent Pelechano, Petra Jakob, Lars M. Steinmetz, Thorsten Zenz, Wolfgang Huber

## Abstract

**Background:** Genome sequencing studies of chronic lympoid leukemia (CLL) have provided a comprehensive overview of recurrent somatic mutations in coding genes. One of the most intriguing discoveries has been the prevalence of mutations in the HEAT-repeat domain of the splicing factor *SF3B1*. A frequently observed variant is predicted to cause the substitution of a lysine with a glutamic acid at position 700 of the protein (K700E). However, the molecular consequences of the mutations are largely unknown.

**Results:** To start exploring this question, we sequenced the transcriptomes of six samples: four samples of CLL tumour cells, of which two contained the K700E mutation in *SF3B1*, and CD19 positive cells from two healthy donors. We identified 41 genes that showed differential usage of exons statistically associated with the mutated status of *SF3B1* (false discovery rate of 10%). These genes were enriched in pathways related to interferon signaling and mRNA splicing.

Among these genes, we found *UQCC* and *RPL31*; notably, a similar effect on these genes was described in a previously published study of uveal melanoma. In addition, while this manuscript was under revision, another study independently reported the common splicing signature of the gene *UQCC* in different tumour types with mutations in *SF3B1*.

**Conclusions:** Our results suggest common effects of isoform deregulation in the genes *UQCC* and *RPL31* upon mutations in *SF3B1*. Additionally, our data provide a candidate list of potential isoform consequences of the SF3B1 (K700E) mutation in CLL, some of which might contribute to the tumourigenesis.

Validation studies on larger cohorts and model systems are required to extend these findings.

## Introduction

Several DNA sequencing studies of chronic lymphocytic leukemia (CLL) revealed that the splicing factor *SF3B1* accumulated somatic point mutations in about 10% of patients [1–3]. In most cases the mutations were located in the genomic regions coding for the C-terminal HEAT-repeat domain and in many cases, the mutations gave rise to specific amino acid substitutions. For instance, the substitution of a lysine to a glutamic acid at amino acid 700 of the protein (K700E) was prevalent in the tumour cells. In addition, the affected amino acids seemed to be clustered spatially in the 3D structure of the protein. These observations suggest that specific changes to the function of the protein could be one of the main drivers of tumour progression in CLL. Additionally, DNA sequencing studies have found recurrent mutations in *SF3B1* in other malignancies, including myelodysplasia (with high incidence in a particular subgroup, RARS) [4, 5] and uveal melanomas [6, 7].

mRNA splicing is the process by which introns are removed from pre-mRNA molecules in order to produce fully mature transcripts. A crucial step of the splicing process is the recruitment of the U2 small nucleolar ribonucleic particle (U2 snRNP) to the branch point sequence: this results in base pairing between U2 snRNP and the pre-mRNA that allows the first chemical reaction of splicing to occur [8]. This recruitment is preceded by the binding of the protein A2AF to the pyrimidine tract and subsequent recruitment of SF3b 155 (the protein encoded by *SF3B1)* [9, 10]. In fact, either blocking the interaction of SF3b 155 to the pre-mRNA sequences using the anti-tumour drug spliceostatin A (SSA) or the knockdown of *SF3B1* results in unstable recruitment of U2 snRNP, which leads to changes in splicing [11].

Additional studies have shown the relevance of *SF3B1* in the regulation of splicing in different biological contexts. For instance, it has been shown that the interaction between SF3b 155 and the proteins Xfp144 and Rnf2 from the Polycomb group of genes is required for the repression of Hox genes during mouse development [12]. In a similar manner, it was shown that the loss of interaction between the proteins coded by the genes *PQBP1* and *SF3B1* alters alternative splicing in mouse neurons and leads to neurite outgrowth defects [10]. These lines of evidence suggest that *SF3B1* is necessary for the correct splicing of pre-mRNAs.

The presence of mutations in *SF3B1* is correlated with adverse prognosis and shorter survival of CLL patients [13]. But despite their usefulness as clinical markers, the functional consequences of the mutations in *SF3B1* are presently not well understood. It has been hypothesized that the mutations in the HEAT-repeat domain might affect the interaction of SF3b 155 with other co-factors and thus, splicing fidelity. Consistent with that hypothesis, it has been observed that mutations in *SF3B1* are associated with the activation of abnormal 3’ acceptor sites of specific genes in CLL tumour cells [1]. In a similar manner, transcriptome analyses of myelodysplastic syndromes and uveal melanomas have identified sets of genes with differential exon usage between tumours with mutations in *SF3B1* and tumours with no mutations in this gene [7, 14].

Here, we aimed to start addressing the potential consequences on isoform regulation of the mutation predicted to cause the K700E substitution in the protein coded by *SF3B1* in CLL tumour cells. We generated transcriptome data from cells of two tumours with mutations in *SF3B1*, two tumours without mutations in *SF3B1* and from cells from two healthy donors. We identified differences in isoform regulation that were associated with the *SF3B1* mutation in 41 genes. We compare our results to previous studies of myelodysplastic syndromes and uveal melanomas with mutations in *SF3B1*.

## Results and Discussion

### Transcriptome-wide data reveal potential isoform regulation associated with mutations in *SF3B1* in CLL tumour cells

We isolated RNA from B-CLL cells of four patients, of which two contained mutations in the *SF3B1* gene (predicted to lead to the K700E substitution in the protein), and two had no mutation in *SF3B1*. Mutation status of each patient was determined by 454 pyrosequencing. We extracted RNA from CD19-purified cells isolated from the peripheral blood of two healthy donors and prepared cDNA libraries for sequencing (see Table S1 for detailed information regarding the samples). We used Illumina HiSeq 2000 to sequence 50 nt paired-end reads using a strand-specific protocol and obtained a total of 275,000,664 sequenced fragments. We mapped the sequencing reads to the human reference genome (*ENSEMBL* release 68) using *GSNAP* (version 2013-05-09), allowing split alignments for exon-exon junctions [15]. We considered only uniquely mapping fragments for further analysis. To observe the expression of the *SF3B1* alleles, we counted the number of mapped fragments in each sample supporting the evidence for the mutation. Based on this, we estimated that when the mutation was present, around half of the transcripts were transcribed from the variant allele (Figure 1). This estimate was consistent with the variant heterogeneity quantification of the tumours’ DNA, as assessed by 454 genomic sequencing (also around 50%, Table S1).

**Figure 1.**
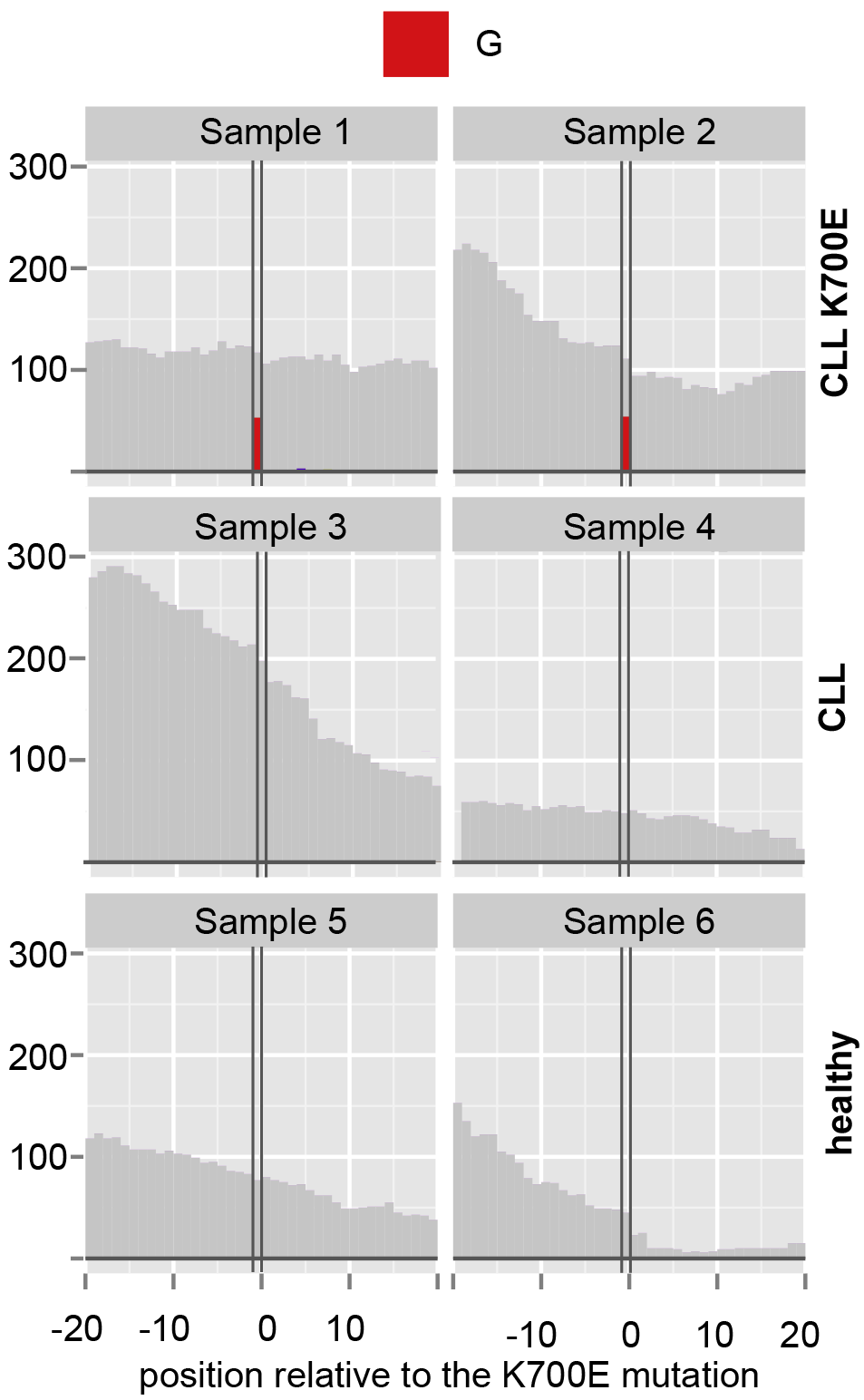
SF3B1 variant expression. Each panel shows the data for one sample. The *y*-axis depicts the genomic coverage resulting from the alignment of the RNA-Seq fragments and the *x*-axis represents the genomic position relative to the position 198,266,834 of chromosome 2 (indicated by the vertical black lines), where the reference genome contains an adenine. The coverage consistent with the reference genome is coloured in gray. The first two samples express the variant that contains a guanine in around half of the transcripts (as indicated by the height of the red bar). This variant is predicted to cause the substitution K700E on the protein coded by *SF3B1*.

We asked, transcriptome-wide, whether specific differences in isoform regulation in the CLL tumour cells were associated with the presence of the *SF3B1* K700E mutation. Therefore, we used *DEXSeq* to test for differences in exon usage (DEU) [16] between the tumour cells with the mutation K700E in *SF3B1* compared to the tumour cells without the mutation and the healthy donors. Briefly, *DEXSeq* considered, for each exon, the ratio between the number of transcripts originating from the gene that contain the exon and the number of all transcripts originating from the gene. This allowed us to identify changes in relative exon usage independently from the fact that a gene could be differentially expressed. Using this approach, we identified a set of 50 exons in 41 genes with DEU at a false disovery rate (FDR) of 10%. This represents an initial candidate list, and analysis of larger cohorts of patients is needed to confirm these associations.

To explore the functions of the genes whose isoform regulation was associated with the mutant *SF3B1* samples, we mapped these genes to pathways annotated in *REACTOME* [17]. We found a statistically significant overrepresentation compared to a background set of genes that were also expressed in these cells, of pathways associated with mRNA splicing and translation at a false discovery rate of 10% (see Table S2). Interestingly, we also found a significant overrepresentation of the interferon signaling pathway, which is known to inhibit cell proliferation and whose aberrant regulation has been linked to aggressive cases of CLL [18]. These results are consistent with the notion that the SF3B1 K700E mutation could preferentially affect the isoform regulation of genes in particular biological pathways.

### The *SF3B1* mutation is associated with differential exon usage patterns seen both in uveal melanoma and CLL

Next we compared our results with those of two previously published transcriptomes. Notably, these studies used the same sequencing technology, had a similar study design (but in different malignancies) and also used the *DEXSeq* method to test for differences in exon usage.

Furney et al. [7] compared three *SF3B1* mutant and nine *SF3B1* wildtype uveal melanoma tumours and identified 34 exons differentially used in 21 genes (10% FDR). Remarkably, we observed significant overlap between their list of differentially spliced genes and those identified in our study (*p* = 2 · 10^−3^, Fisher’s exact test). Specifically, the genes *UQCC* and *RPL31* overlapped with our hits. Furthermore, one out of the two 3’ untranslated regions with DEU that Furney et al. reported for *RPL31*, a gene coding for a ribosomal protein belonging to the 60S ribosomal complex, was also seen as differentially used in our data (Figure 2). This, in principle, could have consequences for the localisation, stability or folding of the RNAs from this locus. Additionally, three out of the four exonic regions that we detected as significant for the chaperone *UQCC* were also detected to be differentially used in the uveal melanoma study. Its authors reported a decrease in the expression of the 3’ end of this gene in uveal melanomas with mutated *SF3B1*, and we observed the same in the CLL tumour cells with mutated *SF3B1* (Figure 3). Interestingly, this region partly codes for a chaperone domain that is conserved with yeast, where it appears to be required for the assembly of the protein ubiquinol-cytochrome C reductase [19]. In humans, genome-wide association studies have linked this gene to body growth [20].

**Figure 2.**
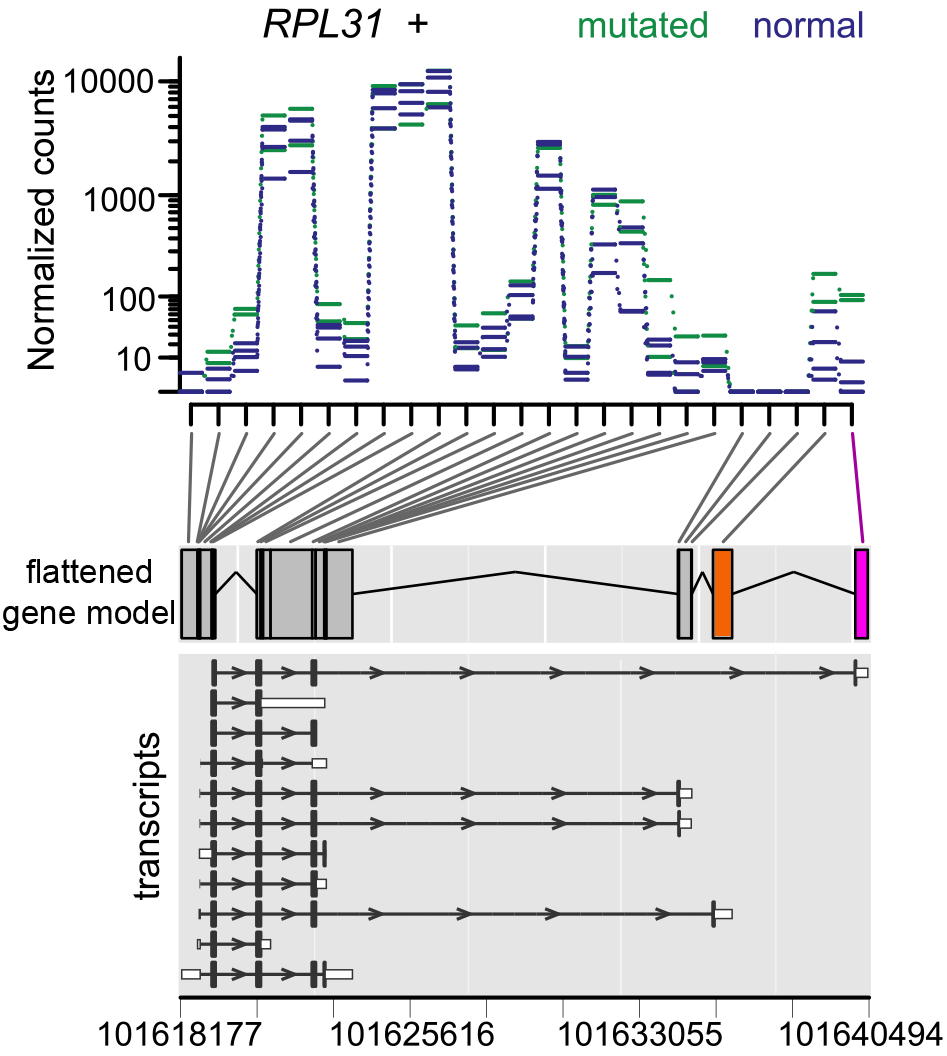
Differential exon usage of gene RPL31. The upper panel shows each exon region of the gene represented along the *x*-axis. The *y*-axis shows the counts for each sample normalised for sequencing depth. The lines from the samples with the mutation in SF3B1 are coloured in green, and the values from the wild-type samples are coloured in blue. The middle panel shows the flattened gene model (set of non-overlapping exon regions) along the genome (*x*-axis). This flattened gene model was derived from the ENSEMBL annotated transcripts (lowest panel) as described in [16]. The exon detected to be significant for DEU in both our study (CLL) and in the uveal melanoma study [7] is coloured in magenta. The exon detected as significant only by the uveal melanoma study is coloured in orange. These exons correspond to 3’ untranslated regions of transcripts (see lower panel).

**Figure 3.**
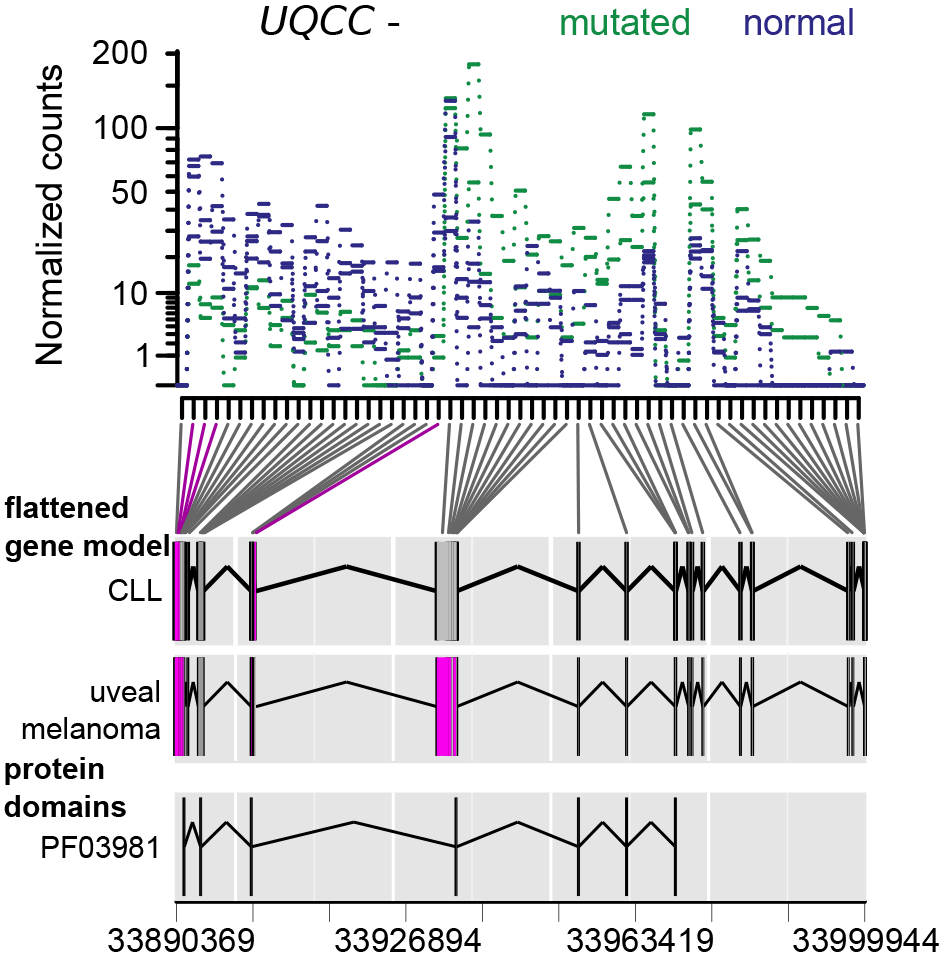
Differential exon usage of gene UQCC. The upper panel shows each exon region of the gene represented along the *x*-axis. The *y*-axis shows the counts for each sample normalised for sequencing depth. The lines from the samples with the mutation in *SF3B1* are coloured in green, and the values from the wild-type samples are coloured in blue. The middle panels depicts two copies of the flattened model, as defined in [16], derived from the transcripts annotated in *ENSEMBL*. In the upper flattened model, the exon regions detected to have significant DEU in our study are coloured in magenta. In the lower flattened model, the significant exon regions from the uveal melanoma study are coloured in magenta. The lowest panel presents the genomic regions coding for the protein domain *PF03981* annotated in *PFAM* [27]. This domain is a highly conserved region of the protein Ubiquinol-cytochrome C chaperone.

Even though we are considering two rather different tumours, and the analyses were done in different laboratories, similar patterns of alternative isoform regulation of the *UQCC* and *RPL31* genes were found to be associated with the mutated status of SF3B1 in both cases. We estimated that the probability of finding by chance that these same exon regions are differentially used was low (*p* = 1.4 · 10^−11^). Hence, our results suggest a link between the mutations in *SF3B1* and the differential exon usage of these two genes that holds across tumour types. Additionally, while this manuscript was under revision, an independent publication validated (using RT-qPCR) the common transcript isoform signature of *UQCC* upon *SF3B1* mutations in both tumour types [21].

Visconte et al. reported 423 exons in 350 genes to be differentially used at an FDR of 5% between myelodysplasia patients with mutations in *SF3B1* and one healthy donor [14]. However, the overlap of their list of genes with our list of genes was not larger than what would be expected by chance, and no common pattern was apparent.

## Conclusion

As a data set relevant to the study of the effects on isoform regulation of the expression of mutant *SF3B1* in CLL, we report transcriptome data from two CLL patients harboring the K700E mutation in SF3B1, two CLL patients without the mutation, as well as two healthy donor cells. Our results provide an initial list of DEU events that appear associated with the K700E mutation in *SF3B1* in CLL (see Supporting Dataset S1). Our data rely on a very limited sample of tumours; substantially larger cohorts (e. g. tens or hundreds of tumours with and without the mutation) will be needed for a more reliable, more comprehensive list of events. A notable result of our analysis is the overlap of events seen here with those in a previous study of uveal melanomas [7], namely, differences in the usage of specific exonic regions of the genes *UQCC* and *RPL31*. This effect was further confirmed for the gene *UQCC* by another independent study using RT-qPCR [21]. These effects could be a prevalent consequence of the mutations in the HEAT-repeat domain of *SF3B1*. The question of whether or not these effects play a causal role in tumorigenesis is not addressed by our analysis, but may merit further study.

## Methods

### Ethics Statement

Samples were acquired by informed written consent in accordance with the Declaration of Helsinki. Ethical and Institutional Board Review (IRB) approvals were obtained from the University Hospital of Heidelberg.

### Sample preparation

Peripheral blood samples from four patients matching standard diagnostic criteria for CLL and featuring a high lymphocyte percentage (median: 98%) were obtained from the University Hospital of Heidelberg. Mononuclear cells (MNCs) were isolated by centrifugation over Ficoll-Paque Premium (GE healthcare, Freiburg). MNCs from buffy coats of healthy donors obtained from the blood bank of the University Hospital of Heidelberg were further CD19-purified by magnetic activated cell sorting (MACS) according to the manufacturer’s instructions (Miltenyi Biotech, Bergisch Gladbach) resulting in purities of ≥ 95% CD19+ cells. Clinical and laboratory data are summarized in Supporting Table S1. Exons of *SF3B1*, *TP53*, *BRAF*, *MYD88* and *NOTCH1* containing mutation hot spots were amplified and subjected to next-generation sequencing on the GS Junior 454 platform (Roche, Penzberg) as in [22]. In brief, genomic regions of interest were amplified from 30 ng genomic DNA in two multiplex-PCRs with 11 primer pairs each. In the same reaction, linker tails were added for ligation of multiplex identifiers and 454-specific adaptors by a second PCR step on the combined pools. Primer sequences are available [22]. Bidirectional sequencing was performed using an emPCR Lib-A kit according to manufacturer’s instructions with adaptations. Sequencing data were processed with GSRunProcessor (v.2.5/v.2.7), performing image and signal processing via the amplicon pipeline (Roche). Variants were identified with Amplicon Variant Analyzer (v.2.5/v.2.7) and annotated manually according to cDNA references from ENSEMBL Genome Browser. Mutations in *SF3B1* were validated by conventional Sanger sequencing and were confirmed to be somatic mutations using whole exome sequencing.

### Strand-specific RNA-Seq library preparation

Total RNA was isolated from 1 · 10^8^ to 5 · 10^8^ cells (depending on the sample) via standard trizol extraction. Strand specific RNA-Seq libraries were prepared as described in [23]. Briefly, polyadenylated RNA was isolated from 10 *µ*g of total RNA using Dynabeads Oligo (dT)25 (Invitrogen) according to the manufacturer’s protocol. The poly(A) enriched RNA was fragmented by incubating the samples at 80^°^C for 4 minutes in the presence of RNA fragmentation buffer (40 mM Trisacetate, pH 8.1, 100 mM KOAc, 30 mM MgOAc). The fragmented RNA was purified using 1.8X (v/v) Ampure XP Beads (Beckman Coulter Genomics) and eluted in 25 *µ*l Elution Buffer (EB) (10 mM Tris-HCl, pH 8) according to manufacturer’s protocol. 24 *µ*l of eluted RNA was reverse transcribed using 1 *µ*l of random hexamers (30 ng/*µ*l, Invitrogen). The samples were denatured at 70^°^C for 5 minutes and transferred to ice. Two *µ*l dNTPs (10 mM), 8 *µ*l 5X first strand buffer (Invitrogen), 4 *µ*l DTT (0.1 M), 0.5 *µ*l actinomycin D (1.25 mg/*µ*l) and 0.5 *µ*l RNaseOut (40 U/*µ*l, Invitrogen) were added to each sample, and the samples were then incubated at 25^°^C for 2 minutes. Following this, 0.5 *µ*l Superscript III reverse transcriptase (200 U/*µ*l, Invitrogen) was added. The retrotranscription was carried out at 25^°^C for 10 minutes, at 55^°^C for 60 minutes, and inactivated at 75^°^C for 15 minutes. The samples were purified using 1.8X of Ampure XP beads and eluted in 20 *µ*l EB. For producing the second cDNA strand, 19 *µ*l of sample was mixed with 2.5 *µ*l of 10x NEBNext Second Strand Synthesis (dNTP-free) Reaction buffer (NEB), 1.5 *µ*l of dNTPs (containing dUTPs instead of dTTPs, 10 mM), 0.5 *µ*l of RNaseH (10,000 U/ml) and 0.5 *µ*l of E.coli DNA polymerase I (10 U/*µ*l, Fermentas). The samples were incubated at 16^°^C for 2.5 hours, 80^°^C for 20 minutes and purified with 1.8X Ampure XP beads, and eluted in 17 *µ*l EB. Two *µ*l end repair buffer and 1 *µ*l end repair enzyme mix (NEBNext DNA Sample Prep Master Mix Set 1, NEB) were added, and the samples were incubated at 20^°^C for 30 minutes. The samples were purified using 1.8x Ampure XP and resuspended in 17 *µ*l EB. Two *µ*l dA tailing buffer (10X NEBuffer 2 from NEB and 0.2 mM dATP) and 1 *µ*l Klenow Fragment 3’-5’ exonuclease (5 U/*µ*l, NEB) were added and the samples incubated at 37^°^C for 30 minutes. The samples were purified using 1.8x Ampure XP and resuspended in 20 *µ*l EB. 2.5 *µ*l 10X T4 DNA ligase buffer (NEB), 0.5 *µ*l multiplexed PE Illumina adaptors (7 *µ*M, Supporting Table S3) and 2 *µ*l T4 DNA ligase were added (2000 U/*µ*l, NEB) and incubated at 16^°^C for 1 h. The dUTPs of the second strand were hydrolyzed by incubating the samples at 37^°^C for 15 min with 1 *µ*l USER enzyme (1 U/*µ*l, NEB) and 5 minutes at 95^°^C. The samples were purified using 0.9 X Ampure XP beads and eluted in 11 *µ*l EB. Enrichment PCR was performed using 5 *µ*l of sample, 25 *µ*l Phusion Master Mix 2x (NEB), 0.5 *µ*l each of oligos PE1.0 and PE2.0 (10 *µ*M, Illumina) and water up to 50 *µ*l final. The PCR program was 30 seconds at 98^°^C, 15 cycles of (10 seconds at 98^°^C, 30 seconds at 65^°^C and 30 seconds at 72^°^C) and 5 minutes at 72^°^C. The PCR product was size-selected (average of 290 bp), and the libraries were submitted for Illumina sequencing.

### Bioinformatics

We mapped the read fragments to the human reference genome from *ENSEMBL* (release 68) using GSNAP (version 2013-05-09) [15, 24]. For each sample, we tabulated the number of uniquely aligned fragments that overlapped with exon annotations from *ENSEMBL* release 68 using scripts based on the python HTSeq library [25]. We used the generalized linear model framework implemented in *DEXSeq* version 1.9.1 to test for differences in exon usage between the samples containing the mutations in *SF3B1* and the wild type allele samples [16].

To avoid biases associated with gene expression strength in further enrichment analysis, we generated a background set of genes that contained at least 600 sequenced fragment counts. We mapped the *ENSEMBL* gene identifiers to pathways annotated in *REACTOME* [17] and tested for overrepresentation of our hits compared to the background using Fisher’s exact test. We corrected for multiple testing using the method of Benjamini and Hochberg [26]. We used the *ENSEMBL Perl API* to convert protein domain coordinates annotated in *PFAM* to genomic coordinates [27]. Genomic ranges operations were performed using the Bioconductor package *GenomicRanges* [28], and visualizations of the genomic ranges were done using *ggbio* [29]. We visualized the coverage vectors and the expression of variants of *SF3B1* using the Bioconductor package *h5vc* [30]. We provide Supporting File S1 with a documented *R* session with the code that was used to analyse the RNA-Seq data and to produce the figures.

## Availability of supporting data

The RNA count data are available in the ArrayExpress database (www.ebi.ac.uk/arrayexpress) under accession number E-MTAB-2025 and in the Bioconductor data package *CLL.SF3B1*.

## List of abbreviations

DEU - differential exon usage; CLL - chronic lympoid leukemia; FDR - false disovery rate

## Author’s contributions

WH, TZ and LMS designed the research. CB, VP, PJ and AR performed the research. TZ and CB selected the patients, isolated the biological samples and performed the mutational analysis. LMS, PJ and VP generated the RNA libraries for sequencing. AR and WH analysed the data and wrote the manuscript with input from all the authors. All authors read and approved the final manuscript.

## Additional Files

Table S1. Clinical and laboratory data of the CLL patients studied.

Table S2. Selected pathways enriched among genes with differential exon usage associated with the SF3B1 mutation (FDR 10%).

Table S3. Oligonucleotide sequences used per sample.

Dataset S1. HTML report of the genes with DEU associated with the mutations in SF3B1

File S1. Documented R session with the program code needed to reproduce our analysis of the data and to generate the figures.

## Acknowledgements

We would like to thank EMBL’s Genomics Core Facility for the RNA sequencing service and the Information Technology (IT) Core Facility for provision of computational infrastructure. WH and AR acknowledge funding from the European Commission through the Collaborative Research Project *RADIANT*.

